# Smart Film Impacts Stomatal Sensitivity of Greenhouse Capsicum Through Altered Light

**DOI:** 10.1101/2020.09.22.309427

**Authors:** Chenchen Zhao, Sachin Chavan, Xin He, Meixue Zhou, Christopher I. Cazzonelli, Zhong-Hua Chen, David T. Tissue, Oula Ghannoum

## Abstract

Optical films that alter light transmittance may reduce energy consumption in high-tech greenhouses, but their impact on crop physiology remains unclear. We compared the stomatal responses of capsicum plants grown hydroponically under control glass (70% diffuse light) or smart glass (SG) film ULR-80, which blocked >99% of ultraviolet light and 19% of photosynthetically active radiation (PAR). SG had no significant effects on steady-state (*g_s_*) or maximal (*g_max_*) stomatal conductance. In contrast, SG reduced stomatal pore size and sensitivity to exogenous ABA thereby increasing rates of leaf water loss, guard cell K^+^ and Cl^-^ efflux, and Ca^2+^ influx. The transition between low (100 μmol m^−2^ s^−1^) and high (1500 μmol m^−2^ s^−1^) PAR induced faster stomatal closing and opening rates in SG relative to control plants. The fraction of blue light (0% or 10%) did not affect *g_s_*, but induced stomatal oscillations in SG plants. Increased expression of stomatal closure and photoreceptor genes in epidermal peels of SG plants is consistent with fast stomatal responses to light changes. In conclusion, light intensity was more critical than spectral quality for optimal stomatal responses of capsicum under SG, and re-engineering of the SG should maximize PAR transmission to maintain a better stomatal development.

**Highlights:**

- Capsicum plants grown under SG film exhibit decreased stomatal pore area, higher water loss and reduced ABA-sensitivity.
- SG-grown plants have faster rates of stomatal closing and opening in response to light intensity changes.
- SG increases efflux of K^+^ and Cl^-^ and influx of Ca^2+^ of guard cells.
- SG upregulated the expression of key genes involved in stomatal regulation and light sensing.

## Introduction

Efficient climatic control in protected cropping can be achieved by alterations in greenhouse structures. These include the even-span greenhouse designed for crop cultivation at high latitude (Sethi, 2009), optimal orientation allowing plants to receive more radiation (Xu *et al*., 2015), different greenhouse shapes to improve the ventilation (Katsoulas *et al*., 2006), and building materials utilising a special plastic film to block UV radiation and enhance light diffusion (Hemming *et al*., 2004). Other techniques such as vent, fog, fan cooling systems, dehumidification, and regeneration process of liquid desiccant also improve glasshouse climatic control (Lefers *et al*., 2016; Rabbi *et al*., 2019; Samaranayake *et al*., 2020; White, 2014). However, the high cost of these solutions indicates that an innovative alternative technique of using low emissivity ‘smart glass’ film, should significantly reduce the costs while maintaining adequate climate control in glasshouses (Lin *et al*., 2020).

The special glass film materials are optically engineered in a nanometre-scale, adjusting light transmittance to allow for high potential of reducing energy cost in high technology greenhouses (Lin *et al*., 2020). The “smart glass” (SG) film ULR-80 blocks the majority of UV light and a proportion of far-red and red light, which can reduce energy load required for heating and cooling in a protected cropping situation (Chavan *et al*., 2020). However, reducing the photosynthetically active radiation (PAR) potentially decreases the growth and productivity of horticultural crops. In a recent study using eggplant grown in a high-tech glasshouse, the application of the SG film led to a net reduction in heat load, water and nutrient consumption and therefore improved energy and resource use efficiency. However, the 19% decrease in PAR reduced fruit yield of eggplants under SG glass by 25% compared to normal control glass (Chavan *et al*., 2020). Whilst SG consistently reduced photosynthetic rates, the response of stomatal conductance was less consistent, decreasing in one season and remaining unaffected in another season (Chavan *et al*., 2020). Optimal stomatal function is crucial for plant photosynthesis (Farquhar and Sharkey, 1982) and water use efficiency (Lawson and Vialet□Chabrand, 2019), but can be compromised under adverse light conditions (O’Carrigan *et al*., 2014). To what extent the effects of altered light conditions generated by a SG film have on stomatal morphology and physiology remains unclear.

PAR, including wavelengths between 400 to 700 nm, supplies the essential photons utilised by plants during photosynthesis, which is highly dependent on its intensity (McCree, 1981). Light directly and indirectly (via photosynthesis) regulates stomatal function (Assmann and Jegla, 2016). Plants have developed sensing mechanisms for both light quantity and quality (Aasamaa and Sõber, 2011; Ballard *et al*., 2019; Düring and Harst, 2015), to adjust stomatal aperture, allowing CO_2_ absorption for carbon fixation. Light also plays important roles in stomatal formation, as well as closing and opening of the guard cells (Roelfsema and Hedrich, 2005). As highly specialized cells, guard cells that form the stomatal pore mediate physiological trade-offs to minimize water loss while maximizing carbon gain in the light. An important limitation in this process is the rate at which stomata open in the light or close in darkness, referred to as stomatal conductance (Drake *et al*., 2013). Rapid stomatal responses to light help to optimise plant photosynthesis (Lawson and Vialet-Chabrand, 2019). Photosynthetic capacity is well linked with the theoretically maximum stomatal conductance (*g_max_*) and operational stomatal conductance (*g_op_*) calculated from stomatal morphological parameters, such as stomatal sizes and stomatal density (McElwain *et al*., 2016).

Long-term effects of light quantity and quality on stomatal density and conductance have been well studied (Savvides *et al*., 2012). Stomatal density increases under high light (Gay and Hurd, 1975), leading to increased stomatal conductance and CO_2_ assimilation (Baroli *et al*., 2008). High light stimulate a rapid stomatal closure along with a rapid production of reactive oxygen species (ROS) (Devireddy *et al*., 2018). ROS accumulation in guard cells activates key ion channels such as slow anion channel (SLAC1) and outward rectifying K^+^ channel (GORK) for stomatal closure (Brandt *et al*., 2012; Deger *et al*., 2015; Lind *et al*., 2015; Zhao *et al*., 2018). Moreover, photoreceptors are key players in the response of plant growth and yield to changes of light environment (Babla *et al*., 2019; Casal, 2013). Blue light-induced stomatal opening is mediated by the light receptor phototropins (PHOT1 and PHOT2) and cryptochromes (CRY1 and CRY2) (Wang *et al*., 2010), while red light induced stomatal opening is mediated by phytochromes (PHYs) (Wang *et al*., 2010). Other light-related genes such as *UV-B Photoreceptor 8* (*UVR8*), *Light-Harvesting Component B* (*LHCB*), and *Ribulose Bisphosphate Carboxylase Small Chain 1* (*RBCS1*) can regulate plant photosynthetic rates (Baroli *et al*., 2008; Borkowska, 2005; Davey *et al*., 2012; Tossi *et al*., 2014; Wang *et al*., 2010; Xu *et al*., 2012). Their responses to SG may elucidate the potential mechanisms that control the stomatal regulation in capsicum.

Our overarching hypothesis was that altered light conditions under SG reduce stomatal density and aperture and affect stomatal sensitivity and guard cell ion fluxes due to regulation of ABA and photoreceptors signalling networks. To address this hypothesis, we used *Capsicum annuum* L., for studying stomatal morphology and physiology. Capsicum, also known as sweet pepper, is the second most cultivated crop after tomatoes in protected cropping in many countries including Australia. Studies on capsicum have mainly focussed on developmental responses to temperature, humidity, and water stress (Bakker, 1989a, b; Hawa, 2003). In this study, we cultivated capsicum plants for 8 months with and without SG film and measured stomatal density, size, guard cell ion fluxes, rate of stomatal response to exogenously applied ABA, and the expression of genes involved in ABA and light signalling networks. We next tested whether signalling pathways triggered by light transitions were altered in a manner that would affect stomatal regulation. We demonstrate that the SG-induced reduction in PAR altered stomatal morphology, behaviour and downstream signalling cascades.

## Material and Methods

### Plant growth and experimental design

The experiment was conducted from April (Autumn in Australia) to Dec 2019 (Summer in Australia) in the state-of-the-art glasshouse facility at Western Sydney University (33°S 150°E, Hawkesbury Campus, Richmond, NSW, 2753 Australia). A detailed description of the facility, including software and climate control is presented by Chavan et al. (2020) and Samaranayake et al. (2020). We used four research bays (105m^2^ each) with precise environmental control of atmospheric CO_2_, air temperature, RH, and hydroponic nutrient and water delivery. *Capsicum annuum* L. seeds (variety Ghia, Syngenta, Australia) were grown in a nursery centre (Withcott Seedlings, Withcott, QLD, 4352 Australia) for six weeks. The seedlings were transplanted in Rockwool slabs and transferred into two control hazed glass (Control) and two SG (Treatment) bays.

The control bays were fitted with HD1AR diffuse glass (70% haze) and the treatment bays had HD1AR diffuse glass, but were also coated with ULR-80 window film, known as “Smart Glass” (SG) (Solar Gard, Saint-Gobain Performance Plastics, Sydney, Australia). The SG film ULR-80 has a low thermal emissivity (0.87) which blocks the light that mainly contributes to heat, but transmits most of the wavelengths of light used by plants for growth in the PAR region. According to the manufacturer specifications, SG blocks around 88% light in the infrared (IR) and far-infrared (FIR) region between 780 nm - 2500 nm; and >99% light in the ultraviolet (UV) region between 300 and 400 nm. SG blocks 43% of total solar energy with 40% transmission, 54% absorption and 6% reflectance. The two control research bays consist of roof glass (70% diffuse light) and wall glass (5% diffuse light). Each bay had 6 gutters with length at 10.8 m and width at 25 cm (AIS Greenworks, Castle Hill, NSW, Australia), which were fitted with 10 Rockwool slabs (90 × 15 × 10 cm, Grodan, The Netherlands) per gutter. Three plants per slab were planted in the four middle gutters, and two plants per slab were planted in the two side gutters which served as buffer plants. Plants were grown in natural light and photoperiod conditions, 25/20°C (day/night) air temperature, 70/80% (day/night) relative humidity, and non-limiting nutrient and water (fertigation) supplied at industry standards. For sample collection and stomatal morphological measurement, unless clarified, top canopy leaves fully exposed to light from each two bays were investigated. Sample collections were completed during sunny conditions on the same day or continuous days to minimise weather effect.

### Relative water loss measurement

Top canopy capsicum leaves which were fully exposed to natural glasshouse light were investigated for relative water loss rate (RWL). RWL was measured using the following equation with modifications (Weatherley, 1950), RWL = (FM – FM_t_)/FM × 100%. Fresh Mass (FM) was determined immediately after samples were collected, and the samples were weighed and recorded as FM_t_ on a scale every ten min for 90 mins. Overall, 10-time points (including 0 min as control) were recorded and the ratio was used to determine differences in the rate of water loss from plants in SG and Control. Five independent leaves from the top canopy of five independent capsicum plants from two bays were collected around 9:00 am on the same day for RWL investigations.

### Stomatal Assay

Stomatal aperture was measured using capsicum epidermal peels from full-expanded top canopy leaves, according to O’Carrigan *et al*., (2014). Epidermal peels were attached to 35-mm glass bottom petri dishes (MatTek Corporation, MA, USA) using silicone adhesive (B-521, Factor II, InC Lakeside, AZ, USA) and bathed in maintaining solution, referred as ‘MS’ (50 mM KCl, 5 mM MES at pH 6.1 with KOH) for about 20 min. Afterward, epidermal peels were imaged in MS under a Nikon microscope attached with a camera and a DS-U3 controller (Nikon, Tokyo, Japan). Images of stomatal apertures and sizes were measured and processed with ImageJ software (National Institute of Health, USA). Stomatal density investigations followed a simplified method (Schlüter *et al*., 2003). Nail polish imprints were taken from the abaxial surface of mature leaves from plants grown under both Control and SG growth rooms. Stomatal densities were determined by light microscopy from leaf imprints of at least five individual plants from both SG and Control growth rooms, respectively. Three independent counts were carried out on each leaf.

The method for measuring ABA-induced stomatal aperture changes followed Cai *et al*., (2017). Manually collected epidermal peels were incubated in MS for 1 h then rinsed with measuring buffer, referred as ‘MB’ [10 mM KCl, 5 mM MES at pH 6.1 with Ca(OH)_2_], three times within 10 min. Epidermal peels were imaged in MB for 10 min under light microscopy as a control. Then, 100 μM ABA treatment was applied and the peels were imaged for another 50 min. Images were taken every 5 min and stomatal apertures were measured and analysed with ImageJ. Thirteen-time points (including 0 min as control) were recorded and the ratio was used to reflect stomatal aperture changes; 30 to 80 stomata from at least three independent epidermal peels were analysed. Epidermal peels were collected at 9 am on sunny days.

### Gas exchange measurement

Leaf gas exchange measurements utilised top canopy leaves which were fully exposed to natural glasshouse light, including net assimilation rate (A_net_) and stomatal conductance (g_s_), were measured using a Li-Cor Li-6400XT infrared gas analyser according Liu *et al*., (2017). For investigations of light intensity on photosynthetic parameters, gas exchange measurements were conducted in three stages and took approximately 140 min. The first stage was established under 1500 μmol m^−2^ s^−1^ PAR for stabilization (20 min, control stage), followed by the second stage when the light intensity was reduced to 100 μmol m^−2^ s^−1^ PAR and maintained for one hour during the measurement. At the initiation of the third stage, the light intensity was returned to 1500 □mol m^−2^ s^−1^ PAR and samples were continuously measured for another one hour. At the control stage, after 20 min measurement, stomatal conductance became stable and the average value was used for calculating relative stomatal conductance to the control stage, which reflects the speed of stomatal movements. For investigations of the blue-light spectrum on stomatal conductance changes, three stages were employed. During the first stage, 1500 μmol m^−2^ s^−1^ PAR [1350 □mol m^−2^ s^−1^ PAR using red LED and 150 μmol m^−2^ s^−1^ PAR using blue LED light (10%)], was employed. After 20 min measurement, 10% blue light was switched off and samples were continuously measured for one hour before the third stage, where 10% blue light ratio was returned, and samples were continuously measured for another one hour. Similarly, average *g*_s_ was calculated and used for normalizing the relative *g*_s_ changes to the control stage. Gas exchange measurements were conducted between 9 am to 3 pm on sunny days and four individual capsicum plants were measured from both SG and Control.

### Stomatal morphological trait measurement and calculation of g_max_

Operating stomatal conductance (*g_op_*) was measured according to Drake *et al*., (2013) and McElwain *et al*., (2016) with modifications using top canopy capsicum leaves. The *g_op_* measurements were taken in the morning on sunny days with licor-6400XT for measurement with 1500 μmol m^−2^ s^−1^ PAR, 70% ambient humidity, 150 μmol m^−2^ s^−1^ air flow and the vapour pressure deficit of ~1 kPa. It is noted that capsicum stomatal opening phase took ~100 min to reach a steady-state *g_op_*. *g_op_* values are means of *g_op_* measurements from four independent capsicum plants using top canopy leaves from both SG and Control at 60-s intervals for the data recording procedures. Maximum theoretical stomatal conductance (*g_max_*) calculation followed Drake *et al*., (2013) and McElwain *et al*., (2016) and utilised stomatal morphological parameters collected based on the stomatal assay:

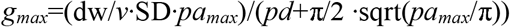

where dw = diffusivity of water vapour at 25 °C (0.000025 m^2^ s^−1^) and *v* = molar volume of air (0.022 m^3^ mol^−1^) are both constantsMcElwain *et al*., (2016), SD is stomatal density (m^−2^) observed from our stomatal assay, stomatal pore sizes (m^2^) were calculated as an elipse using stomatal pore length (m) as the long axis and ½ stomatal pore width (m) as the short axis and *pa_max_* (maximum stomatal pore size) was recorded from each replicate among four independent plants; *pd* is stomatal pore depth (m) considered to be equivalent to the stomatal width of an fully turgid guard cell (McElwain *et al*., 2016).

Similarly, stomatal sizes (μm^2^) were calculated following an elipse using stomatal length (μm) as the long axis and ½ stomatal width (μm) as the short axis, maximum stomatal sizes (*SS_max_*, μm^2^) were recorded accordingly from each replicates among four independent capsicum plants. Stomatal opening and closing half-times were calculated from the gas exchange measurement, where stomatal opening half-time was calculated as the time it took to reach the maximum stomatal conductance in response to the light transition from 100 μmol m^−2^ s^−1^ to 1500 μmol m^−2^ s^−1^ PAR, and stomatal closing half-time was calculated by the time capsicum plants took for reaching the minimum stomatal conductance in response to the light transition from 1500 μmol m^−2^ s^−1^ to 100 μmol m^−2^ s^−1^ PAR. Stomatal opening and closing half-times were recorded from four independent capsicum plants of both SG and Control.

### Guard cell ion fluxes measurement

For guard cell ion flux measurements, the preparation of epidermal peels was identical to the stomatal bioassay. Net fluxes of K^+^, Cl^-^, Ca^2+^, and H^+^ were measured using non-invasive, ion-selective microelectrodes (MIFE) on guard cells of capsicum according to Pornsiriwong *et al*., (2017) and Zhao *et al*., (2019). Specific details related to the MIFE theory, electrode fabrication and calibration are described in Shabala *et al*., (2013). Epidermal peels were pre-treated with MS for 20 min before blue light treatment. The peels were fixed on a coverslip coated with silicone adhesive and then placed in a long, flat 5 mL measuring chamber containing MB. Electrodes with fine tips (Resistance = 4 to 6 GΩ) were filled with ion-selective ionophore cocktails (Sigma, Buchs, Switzerland) and their tips were moved towards and away from the sample in a slow (5 s cycle, 80 μm amplitude) square-wave by a computer-driven micromanipulator. Net fluxes of ions from guard cells were calculated from the measured differences in electrochemical potential for these ions between two positions. Net K^+^, Ca^2+^, H^+^, and Cl^-^ fluxes from guard cells were measured for 10 min as a control to ensure initial, steady values before implementing the blue light treatment and then measurements were conducted for another 30 to 40 min. At least five individual stomatal guard cells from independent plants were investigated for ion flux measurements.

### Quantitative real time-PCR

Quantitative real-time PCR was performed as previously described (Chen *et al*., 2016). We measured the transcripts of key genes of abaxial epidermal peels of capsicum leaves. The details of tested genes can be found in Table S1. Epidermal peels were collected for gene expression investigations to minimize the effect of mesophyll cell mRNA and to enrich the guard cell mRNA (Cai et al., 2017). Fully expanded leaves from the top canopy of four-month-old capsicum plants were selected for sample collection. Under normal light inside growth rooms, the epidermal peel was immediately collected and stored in liquid nitrogen. Total RNA was extracted using a RNeasy Plant Mini Kit (Qiagen, Australia) following the manufacturer’s procedure, and the residual genomic DNA was removed with amplification grade DNase I (Ambion). First-strand cDNA was synthesized with the SensiFAST Kit (Bioline, Alexandria, Australia). Fluorescence reflecting target genes expression was determined by the SensiFAST SYBR No-ROX Kit (Bioline, Australia) using gene-specific primers (Table S1) by employing a Rotor-Gene Q6000 (Qiagen). qPCR conditions were composed of three steps of cycling: polymerase activation at 95 °C for 15 min; 40 cycles were set up for denaturation at 94°C for 15 s, annealing for 15 s at 55 °C, extension at 72 °C for 15 s; SYBR green signal data were acquired at the end. Ubiquitin-conjugating gene (*UBI-3*) (Wan *et al*., 2011) was used as the reference for normalization of relative gene expression. Data were expressed as the average of four independent plants from two research bays with two technical replicates.

### Statistical analysis

Statistical significance between SG and Control plants, before and after treatment was analysed using Student’s t-test and SPSS one-way ANOVA test was applied for statistical analysis of ion flux measurement. All data were presented as means with standard errors.

## Results

### Smart glass (SG) reduced stomatal pore size but not stomatal conductance or density

Light conditions are vital for stomatal formation and development. To investigate stomatal morphological changes induced by SG, we measured stomatal parameters from both control and SG grown plants. Relative to the control glasshouse bays, application of the SG film ULR 80 blocked 99% of UV, 58% of far-red, and 26% of red light, along with a 19% reduction in PAR. The SG chambers appeared light blue and the control chambers appeared white from the aerial view (Fig. S1A-B). Compared with control plants grown under normal glass condition, plants grown under SG had similar stomatal conductance (Fig. 1B). However, SG significantly decreased stomatal pore size (*P* = 0.036) by 13% relative to the control (Fig. 1C), due to reduced stomatal pore length rather than width (Fig. 1A; Table S2). Stomatal size and density were not statistically different between the control and SG treatments (Fig. 1D-E). These results partially support our hypothesis that altered light conditions under SG will reduce stomatal aperture, indicated by decreased stomatal pore size (length) but not stomatal size or density.

**Fig 1.**
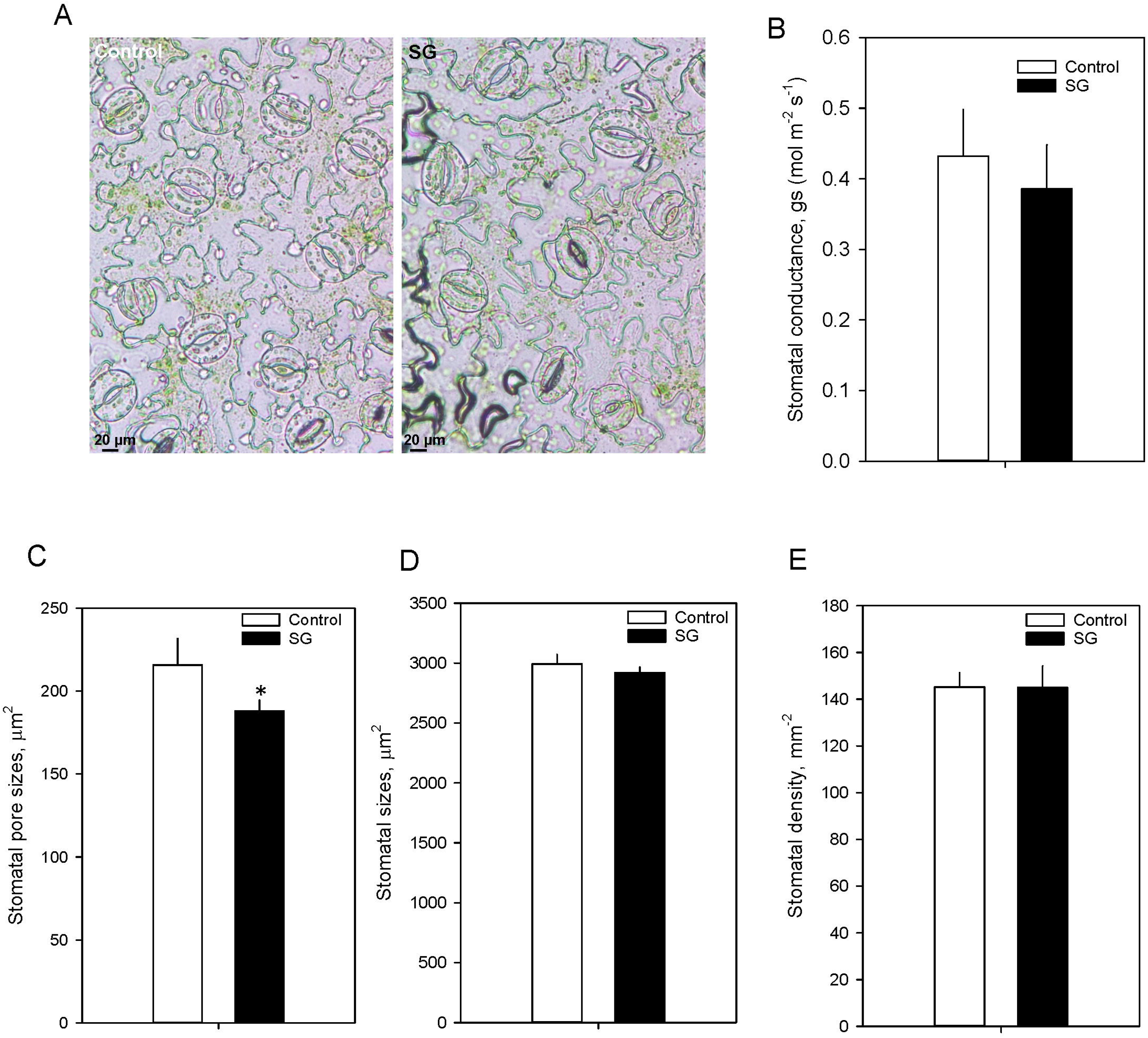
Effect of smart glass (SG) on stomatal traits of capsicum. **(A)** representative stomatal photos collected from epidermal peels of both control and SG. Scale bar = 20 μm **(B)** stomatal conductance in the Control and SG under control conditions (1500 PAR at initial stable stage). **(C-E)** stomatal pore size, stomatal size and stomatal density in the control and SG (n=5 biological replicate with 50-80 stomata). Values are means ± SE. *P<0.05.

### SG led to greater leaf water loss, slower ABA-induced stomatal closure and upregulation of ABA signalling genes

Did changes in stomatal morphology observed under SG induce physiological or molecular changes in the stomatal response? To answer this question, we compared water loss rate between SG and control plants and subsequently investigated genetic transcripts relating to ABA signalling networks. During the initial 40 min following leaf detachment, SG and control plants had similar rates of relative water loss. After 60 min, SG leaves transpired water faster than control leaves (*P* = 0.012 at 90 min) (Fig. 2A). Given stomata mediate the majority of plant water loss, more water loss from leaves of SG plants indicates a change in stomatal responses. Thus, stomatal closure rate in response to ABA was investigated using epidermal peels, and the initial stomatal aperture before ABA application was not significantly different between the control and SG treatments (Fig. 2B). Exogenously applied ABA caused a slower stomatal closure in SG plants, especially after 40 min of incubation with ABA (Fig. 2B).

**Fig 2.**
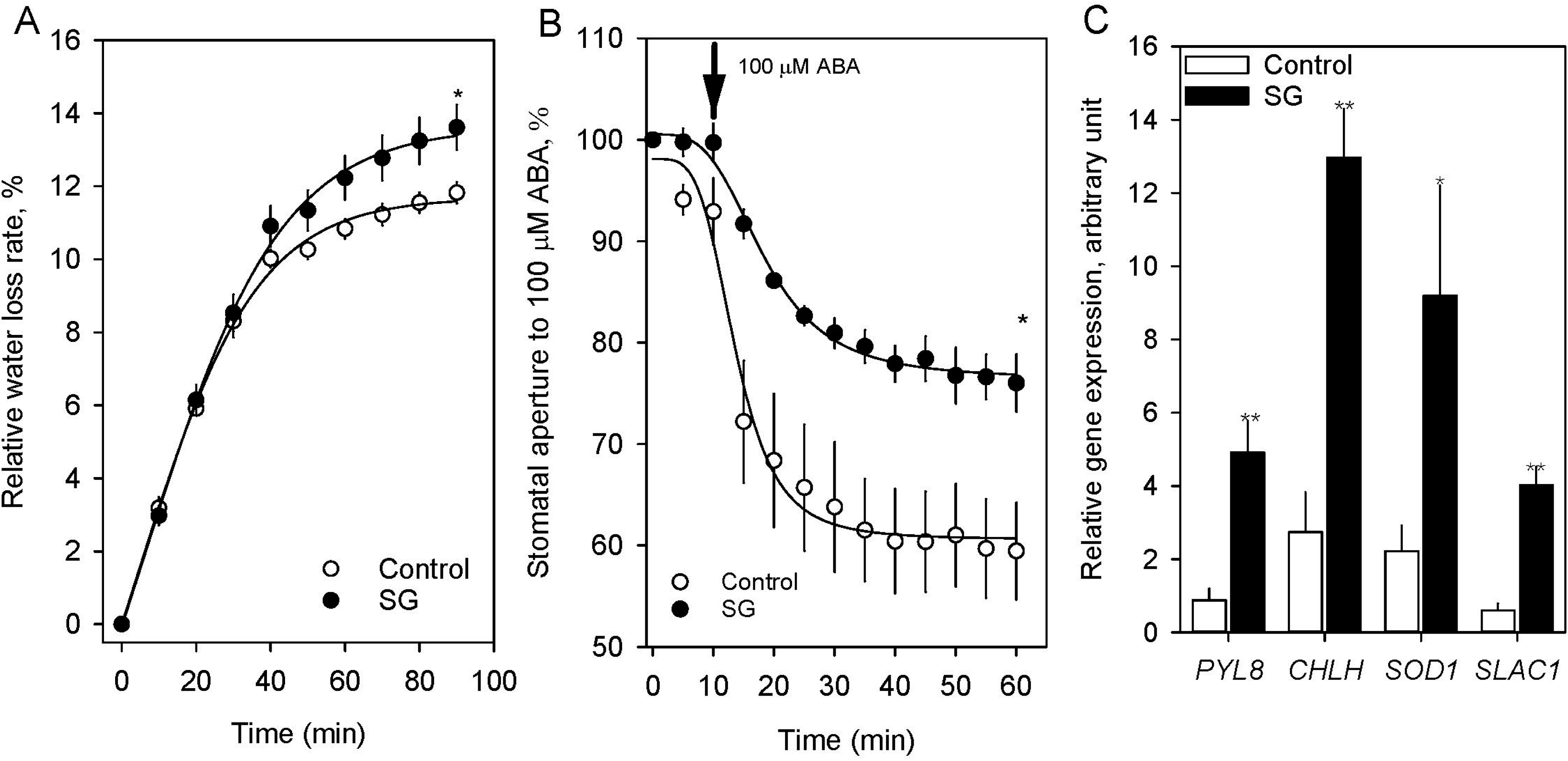
Smart glass (SG) alters relative water loss, ABA-induced stomatal closure, and expression of genes of ABA signalling in capsicum. **(A)** relative water loss was calculated based on the mass loss every ten min. Values are means ± SE (n=5). **(B)** stomatal response to exogenous 100 μM ABA. Data are mean ± SE (n=4 biological replicates from 30 to 80 stomata). **(C)** relative expression of genes of ABA signalling from epidermis. *Pyrabactin resistance8* (*PYL8*), *Mg-chelatase H subunit* (*CHLH*); *superoxide dismutase* (*SOD*) and *slow anion channel-associated 1* (*SLAC1*). Data are means ± SD (n=4 biological replicates with 2 technical replicates). *P<0.05; **P<0.01.

We then quantified the expression of genes involved in ABA signalling in epidermal peels. *PYL8* and *CHLH* are vital ABA receptors whose mutations both lead to severe open stomata and ABA-insensitive phenotype, whilst overexpression of *PYL8* or *CHLH* leads to high degrees of stomatal closure (Gonzalez-Guzman *et al*., 2012; Lim *et al*., 2013; Shen *et al*., 2006). Relative to the control, there were significant increases (~ 4-6-fold) in *PYL8* and *CHLH* transcripts (Fig. 2C). Since ROS accumulation has been identified as a central network component for stomatal closure (Sierla *et al*., 2016), core genes encoding ROS metabolism were also investigated [e.g. SOD catalyses the decomposition of hydrogen peroxide (H_2_O_2_), the GTP binding protein ADP-ribosylation factor 1 (ARF1) (Dana *et al*., 2000)]. SG generated a four-fold upregulation of *SOD*, and *ARF1* expression was enhanced by 5-fold in SG compared to the control (Figs 2C and S4). However, the SG treatment showed no significant effect on the expression of *Catalase 3* (*CAT3*), which catalyses the breakdown of H_2_O_2_ into water and oxygen (Fig. S4). Finally, the expression of *SLAC1*, whose protein contributes to stomatal closure (Deger *et al*., 2015), was four-fold higher in SG compared to control epidermal peels (Fig. 2C). Taken together, these results support our hypothesis that SG will affect stomatal sensitivity to water stress and ABA-mediated signalling processes.

### SG stomata responded faster to light transitions without changes in SS_max_, g_max_ or g_op_

To investigate if the SG treatment has altered stomatal sensitivity, we measured changes in stomatal aperture in response to light transitions from 1500 to 100 μmol m^−2^ s^−1^ PAR (Fig. 3A). On average, stomata of SG plants showed lower opening and closing half-times relative to the control (Fig. 3B). In particular, SG stomata closed faster in response to the transition to low (100 μmol m^−2^ s^−1^) PAR, and opened faster in response to the subsequent transition to high PAR (1500 μmol m^−2^ s^−1^) (Figs 3A-B). After 140 min of both light transitions, stomatal conductance was significantly higher in SG relative to control plants (Fig. 3A, Table S2, P < 0.01).

**Fig 3.**
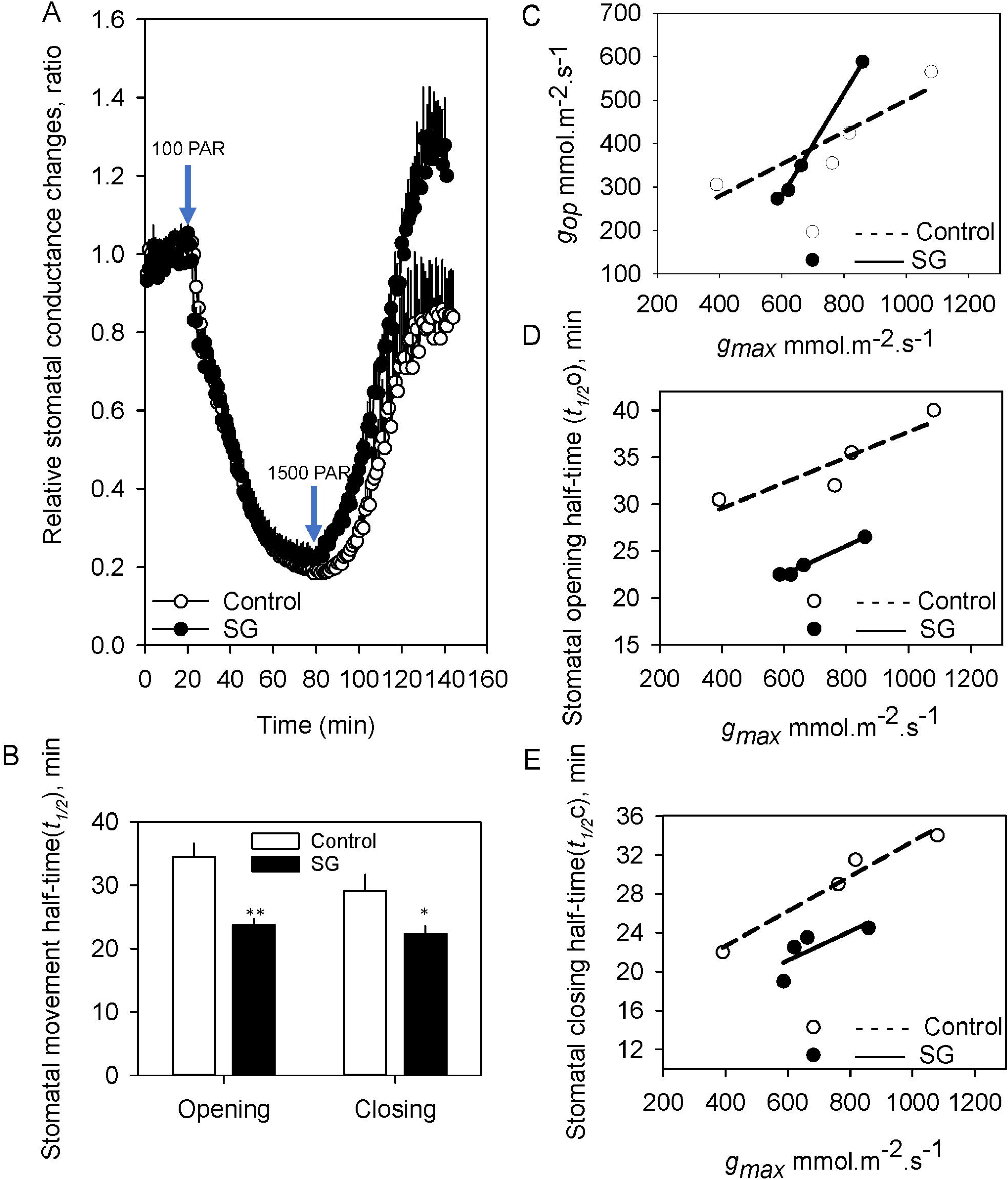
Effect of smart glass (SG) on stomatal sensitivity to light transitions in capsicum. **(A)** Stomatal conductance was initially stabilized and recorded under 1500 PAR then reduced to 100 PAR for 1 hour and followed by 1500 PAR for another 1 h. The averaged stomatal conductance value from the initial 20 min was used for normalizing stomatal conductance ratios to the initial control stage. **(B)** half stomatal opening time (t1/2o) and half stomatal closure time (t1/2c) in response to light transitions. **(C**-E) correlation analysis between maximum theoretical stomatal conductance (*g_max_*) and operational stomatal conductance (*g_op_*) and stomatal opening and closing half-times (*t_1/2_*). Data are means ± SE (n=4). *P<0.05; **P<0.01.

Given the clear link between light conditions and stomatal development (Fu *et al*., 2010; O’Carrigan *et al*., 2014), we correlated stomatal parameters with maximum theoretical stomatal conductance (*g_max_*) and operational stomatal conductance (*g_op_*). SG and control plants maintained similar *g_op_* and *g_max_* (Table S2). The relationship between *g_op_* and *g_max_* was steeper in SG than control plants (Fig. 3C). Both treatments showed parallel relationships between opening and closing half-times with *g_max_* (Fig. 3D-E) and *g_op_* (Fig, S3C-D). The maximal stomatal size (*SS_max_*) also showed similar relations with *g_op_* in both treatments (Fig. S3A-B). Overall, SG produced more active stomata in response to light intensity changes with smaller aperture, but not size, supporting our hypothesis that stomatal sensitivity to light conditions will increase due to the altered light condition under SG.

### SG enhanced expression of photoreceptor and photosynthesis genes in epidermal peels

Stomatal conductance was investigated in response to changes in blue light fraction, which is required for inducing stomatal opening (Inoue and Kinoshita, 2017). SG and control plants responded similarly to blue light (Fig. 4A). During the measurement, there were no differences in stomatal conductance under 1500 μmol m^−2^ s^−1^ PAR (Fig. 4B). Removing 10% blue light (1350 μmol m^−2^ s^−1^ PAR) generally induced stomatal closure in both SG and control leaves, while the retrieval of 10% blue-light increased stomatal conductance similarly in both treatments (Fig. 4B). Accordingly, our hypothesis about increased stomatal sensitivity to blue light under SG is rejected.

**Fig 4.**
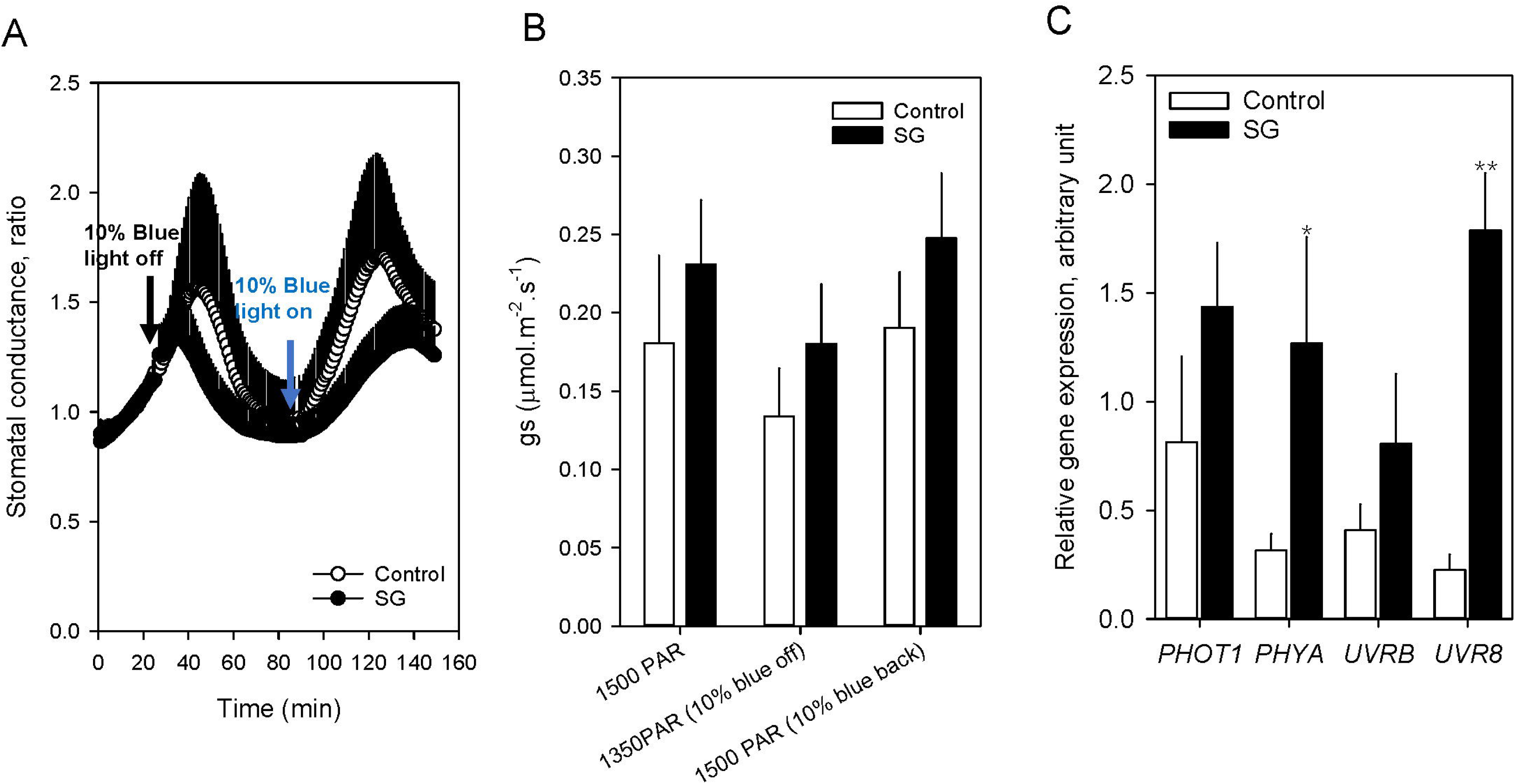
Smart glass (SG) induces different stomatal responses to blue light and gene expression in capsicum. **(A)** stomatal conductance was monitored in three light conditions: normal light (1500 PAR: 1350 Red + 150 Blue) for 20 min, 1350 Red PAR for 1 hour, and normal light for another 1 h. The averaged stomatal conductance value from the initial 20 min measurement was used for normalizing stomatal conductance ratios. **(B)**, bar graphs are three points of the end of each light condition. Data are means ± SE (n=4) **(C)** gene expression of *Phototropin 1* (*PHOT1*), *Phytochrome A* (*PHYA*), *UV response elements* (*UVRB*) and *UV-B Receptor 8* (*UVR8*). Data are means ± SE (n=4 biological replicates with 2 technical replicates). *P<0.05; **P<0.01.

Capsicum leaf epidermal peels were used to assess the expression of photoreceptor and photosynthesis associated genes such as *PHOT1, PHYA*, and *RBCS1*. Compared with the control, SG grown plants had enhanced gene expression by 75% in *PHOT1*, 300% in *PHYA* and 165% in *RBCS1* (Figs 4 C and S4). Further, SG plants exhibited increased gene expression of *UV light response element* (*UVRB*) and *UVR8* relative to control plants (Fig. 4C). These are crucial genes regulating photosynthesis and differential gene expression patterns between SG and control plants, indicating that a different factor was deployed by SG plants to adapt to changed light conditions. Overall, SG stomata maintain a higher sensitivity to light at physiological and molecular levels under glasshouse conditions.

### Higher guard cell flux of K^+^, Ca^2+^, and Cl^-^ is induced by SG but suppressed by blue light

Stomatal opening and closing are regulated by ion fluxes across membranes of guard cells. We investigated ion fluxes from guard cells of both treatments. In normal light, guard cells from SG plants showed approximately three times greater efflux of K^+^ and Cl^-^ compared to control plants (Fig. 5A and D), which reduced stomatal aperture and contributed to closure. The Ca^2+^ influx of guard cells from SG plants was about two times higher than that from control plants (Fig. 5B). The H^+^ efflux of guard cells was similar between SG and control plants under normal light (Fig. 5C).

**Fig 5.**
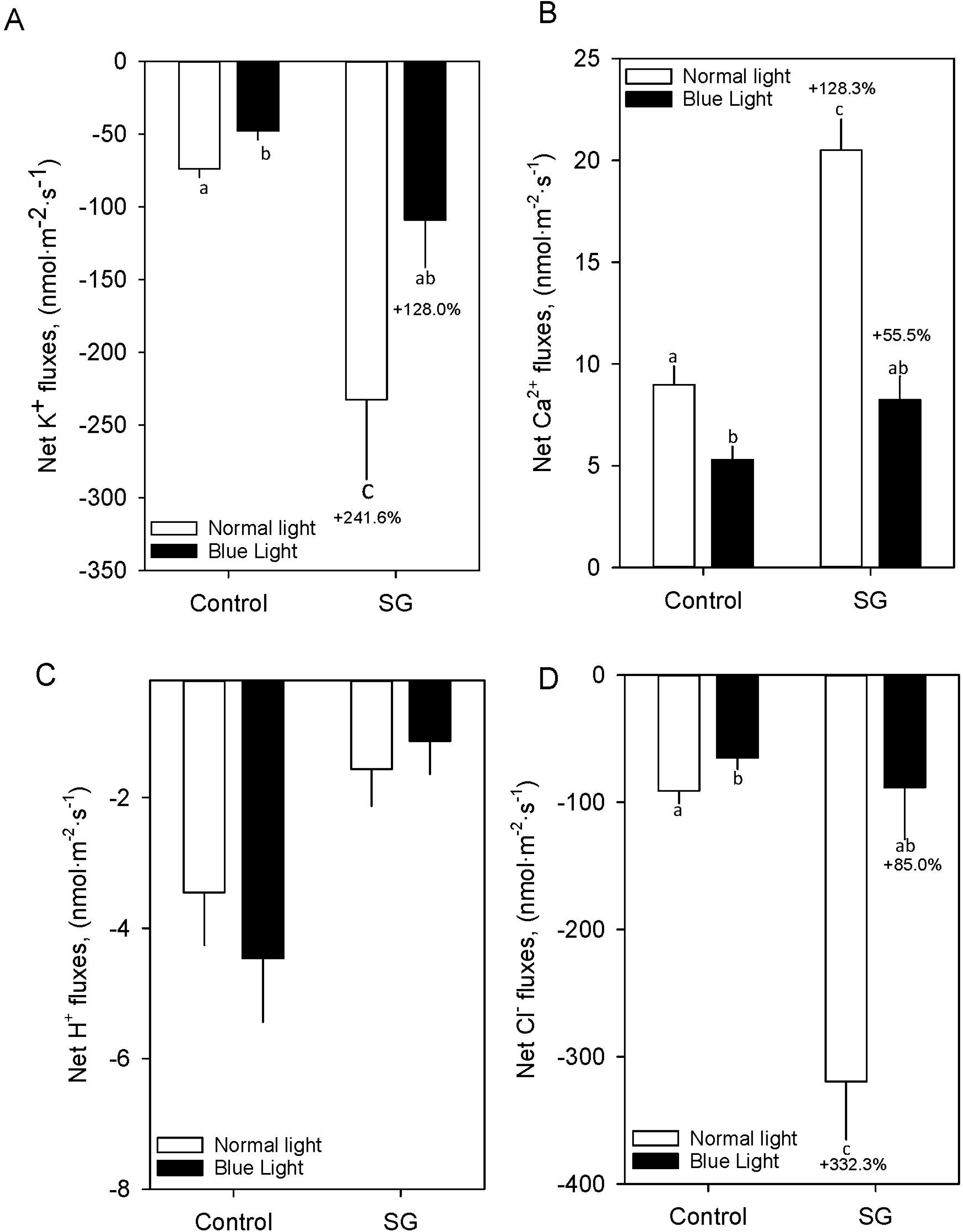
Smart glass (SG) affects ion fluxes and their regulation by blue light in guard cell of capsicum. Net fluxes of K^+^ **(A)**, Ca^2+^ **(B)**, H^+^ **(C)** and Cl^-^ **(D)** were recorded from guard cells in leaf epidermal peels. Data are means ± SE (n=5 to 7 plants). Different lowercase letters represent the statistical difference.

Blue light significantly suppressed K^+^ efflux of guard cells by 35% in control and 53% in SG plants (Fig. 5A). Moreover, blue light also suppressed Cl^-^ efflux by 72% in SG and 28% in control plants (Fig. 5D). Meanwhile, Ca^2+^ influxes were suppressed by 41% and 60% in control and SG, respectively (Fig. 5B). In contrast, blue light slightly induced H^+^ efflux in the control treatment, but slightly suppressed H^+^ efflux in SG, indicating similar effects on SG and control plants guard cells (Fig. 5C). Overall, SG epidermal peels indicated enhanced solutes loss under normal light condition but maintains ability to response to blue light. This agrees with our hypothesis that long-term altered light conditions under SG will affect guard cell ion fluxes determining stomatal status.

## Discussion

In this study, we compared stomatal functions in upper canopy leaves from capsicum plants grown under SG and control lighting environment. Our results can be categorised into four main findings. Firstly, SG reduced stomatal pore size and increased guard cell fluxes (K^+^, Cl^-^ efflux, and Ca^2+^ influx) and the expression levels of *SLAC1* involved mechanism of cellular ion homeostasis without appreciably affecting g_s_, *g_op_*, *g_max_*, stomatal size or density. Secondly, SG reduced stomatal sensitivity to ABA, leading to relatively more water loss in detached leaves, and this response was underpinned by upregulation of ABA (*PYL8* and *CHLH*) and ROS (*SOD1* and *ARF1*) related genes. Thirdly, SG stomata responded faster to PAR transitions, such that stomatal opening and closing speed was proportional to *g_max_*, whilst the relationship between *g_op_* and *g_max_* was steeper in SG plants. Fourthly, even though SG filtered out light most efficiently in the blue spectrum, dependence of stomatal conductance on blue light was similar between SG and control treatments. Yet, guard cell fluxes showed selectively greater (K^+^ and Ca^2+^) or different (H^+^) blue light sensitivities in the SG plants, and this was associated with increased expression of photoreceptor genes (*PHOT1* and *PHYA*) and UV-B light response genes (*UVRB*, *UVR8*). Combining all these findings, SG light condition did not impair stomatal ability to respond to light changes of capsicum leaves; instead, the adaptation of SG capsicum stomata to the altered light condition involved a more active response to PAR changes, ABA signalling, and solutes loss to maintain a decreased stomatal pore size under SG light condition.

### Decreased stomatal pore area in SG is underpinned by enhanced guard cell solute loss and anion channel activity rather than changes in stomatal morphology

Stomata regulates plant water-use efficiency by affecting CO_2_ uptake and photosynthesis as well as transpiration (Brodribb *et al*., 2009). Under low light conditions, where light reception is limited, full stomatal opening may not be necessary for photosynthesis (Pasternak and Wilson, 1973). A study in sweet pepper suggests that partial shade induced lower stomatal aperture (Jaimez and Rada, 2011). Under SG, where the light intensity was lower than in the control, stomatal pore sizes were significantly smaller in SG than control leaves, due to decreased stomatal pore length (Fig. 1C). We found that there was no difference in stomatal density between SG and control plants (Fig. 1E). Moreover, no significant difference was observed in most of the stomatal morphological parameters (Table S1), suggesting that ion flux changes may affect the stomatal aperture.

Stomatal opening requires activation of potassium inward channels, such as KAT1, KAT2 (Ronzier *et al*., 2014), and AKT1 (Nieves-Cordones *et al*., 2012), as well as decreased channel activities of potassium outward channel GORK (Hosy *et al*., 2003). SLAC1 plays a vital role in regulating stomatal response to light (Hiyama *et al*., 2017), CO_2_ (Lind *et al*., 2015), and humidity (Vahisalu *et al*., 2008); stomatal closure in response to drought (Geiger *et al*., 2009), salinity (Qiu *et al*., 2016), and darkness (Merilo *et al*., 2013). Here, SG stomata exhibited significantly higher guard cell efflux of K^+^ and Cl^-^, which suggests that SG plants close stomata more rapidly than control plants (Fig. 5A and D). Compared with control, SG plants take up about twice more Ca^2+^ into guard cells (Fig. 5B), which is in agreement with other studies which showed that increased cytosolic Ca^2+^ activates anion channel (Asano *et al*., 2012), deactivates potassium inward channels (Ronzier *et al*., 2014) for stomatal closure (Asano *et al*., 2012; Zhao *et al*., 2018). In our study, the higher guard cell efflux of K^+^ and Cl^-^ and Ca^2+^ influx under SG reduces cell turgor, thereby decreasing stomatal pore area. SG plants also showed significantly higher expression of *SLAC1* responsible for Cl^-^ efflux (Brandt *et al*., 2012), and higher expression of ABA receptor genes (Fig. 2C). Hence, we propose that SG-induced prolonged low light conditions may increase solute loss, leading to a decrease in stomatal pore area and reduced stomatal conductance.

### SG deceased stomatal sensitivity to exogenously applied ABA due to up-regulated ABA signalling leading to higher water loss from capsicum leaves

Sensing adverse environments and producing ABA for closing stomata has been well established during plant evolution (Lind *et al*., 2015), and the speed for closing stomata reflects the plant’s ability to adapt to a new environment (Pantin *et al*., 2013; Wang and Chen, 2020). A higher relative water loss rate in SG leaves and stomata on plants grown under SG do not open as wide and close more slowly due to the ABA application, indicating that a modified acclimation mechanism for closing stomata developed in plants grown in SG.

We investigated transcripts of the critical components of ABA signalling networks. The ABA-induced signalling network consists of critical components, including ABA receptors (Gonzalez-Guzman *et al*., 2012; Merilo *et al*., 2013), ROS production (An *et al*., 2008), Ca^2+^ signalling (Asano *et al*., 2012; Ronzier *et al*., 2014) and regulation of ion channels (Deger *et al*., 2015; Hosy *et al*., 2003; Vahisalu *et al*., 2008). In our study, SG increased expression of ABA receptor gene *PYL8* and ABA signalling genes *CHLH* and *ARF1* (Figs 2C and S4), indicating their roles are affected by SG (Liu *et al*., 2013; Mishra *et al*., 2006). SOD1 functions as a strong ROS remover in plants, which also affects stomatal activity through ROS accumulation (An *et al*., 2008; Jannat *et al*., 2011; Jiang and Yang, 2009). Therefore, the upregulated gene expression of *SOD1* may suggest ROS accumulation in SG guard cells as part of the SG affected ABA signalling (Figs 2C and S4) to regulate anion channels for stomatal closure (Sierla *et al*., 2016; Zhao *et al*., 2018). This was confirmed by increased expression of *SLAC1* in SG plants, enhanced guard cell Cl^-^ efflux, and the slow stomatal response to exogenous ABA treatment. As the ABA induced stomatal signalling elements were already enhanced, exogenously applied ABA failed to induce further significant stomatal closure in SG plants, leading to a higher water loss rate in SG detached leaves.

### Faster SG-induced stomatal response to light transitions correlates with *g_max_* without affecting photosynthesis rate

Stomatal morphology (e.g. stomatal size), stomatal conductance, and photosynthesis rate are linked to *g_max_* and *g_op_* (Drake *et al*., 2013), such that faster stomatal reaction speed to light increases *g_op_*, thereby improving photosynthesis and water use efficiency (Lawson and Matthews, 2020). We found that SG stomata exhibited faster response rate to light transitions and these rates were linearly correlated with *g_max_*. Further analysis showed that SG plants had a narrow range of *g_max_*, even though averages of *g_max_* and *g_op_* were not affected by SG. Low PAR generally reduces stomatal conductance and net photosynthetic rate (Farquhar and Sharkey, 1982; Pasternak and Wilson, 1973; Roelfsema and Hedrich, 2005). This is also supported by a previous study, where capsicum plants grown in low light conditions (20% of control) had reduced stomatal index and CO_2_-saturated photosynthesis rate (Fu *et al*., 2010). Here, we found that SG plants exhibited a decreased *g_max_* range and produced slightly smaller stomata, which partly agrees with the above findings. However, SG did not significantly affect net photosynthetic rate (Fig. S2), which may be due to a similar *g_op_* range and relatively high *g_max_*.

The ‘smaller but faster stomata’ theory was supported by a multi-species study which found that smaller stomata were usually associated with faster stomatal dynamics (Drake *et al*., 2013; Franks and Beerling, 2009), was summarised in a recent review (Lawson and Vialet□Chabrand, 2019). Evolution of faster stomatal response promoted expansion of grasses (Chen *et al*., 2017), leading to higher plant productivity, efficiency and fitness (Lawson and Vialet□Chabrand, 2019). In our study, SG plants exhibited smaller stomatal pore size as well as faster opening and closing speed in response to light transitions between 1500 μmol m^−2^ s^−1^ PAR to 100 μmol m^−2^ s^−1^ PAR, which validate the ‘smaller but faster stomata’ theory (Drake *et al*., 2013) in a greenhouse horticultural crop.

### Enhanced expression of light responsive genes underpins the effects of SG on capsicum

Photoreceptors are closely linked with plasma membrane transport, determining plant ionic balance, affecting plant growth, development, and yield (Babla *et al*., 2019). PHOT1 and PHOT2 were reported to regulate membrane transport via regulating cytosolic Ca^2+^ in plants (Briggs and Christie, 2002). Further evidence suggests that PHOT1 joins blue light induced Ca^2+^ influx to the cytoplasm and therefore affects significant changes of Ca^2+^ and H^+^ fluxes (Babourina *et al*., 2002). In our study, SG plants showed upregulation of *PHOT1* along with increased Ca^2+^ influx. SG reduces 99% of UV light into the greenhouse bays, which also induced a significantly higher expression of UVR8 (Fig. 4C). In *Arabidopsis*, UVR8 plays important roles in UV light induced stomatal closure by a mechanism involving both H_2_O_2_ and NO generation in guard cells (Tossi *et al*., 2014). This further supports our observations of a higher basal level of expression of light responsive genes in SG shows that stomata maintain full capacity to respond to light alternations.

To confirm previously reported blue light induced stomatal opening case studies, where crucial ion channels, such as K^+^ (Takahashi *et al*., 2013), Ca^2+^ (Ronzier *et al*., 2014), H^+^ (Inoue and Kinoshita, 2017) and Cl^-^ (Hiyama *et al*., 2017), were regulated by blue light, we measured dynamic stomatal conductance during blue light transitions and ion flux changes in response to blue light with no red light background. SG plants exhibited a suppression of K^+^ and Cl^-^ effluxes and Ca^2+^ influx (Fig. 5). Absence of blue light slightly decreased stomatal conductance in both SG and control plants, whilst blue light retrieval mildly increased stomatal conductance (Fig. 4B), indicating the key role of blue light in stomatal opening in capsicum (Inoue and Kinoshita, 2017). However, compared with the light retrieval-induced stomatal dynamic changes (Fig. 3A), SG plants stomata obviously responded more actively to light intensity than blue light spectrum (Figs 3A-B and 4A). Overall, SG plants maintain a similar capacity of responding to blue light spectrum but more actively respond to light intensity relative to the control, and this is highly linked with the increased expression of light responsive genes (Fig. 4C).

## Conclusions

Altered light condition in SG did not lead to strong stomatal morphological changes but decreased stomatal aperture, which is the consequence of vigorously activated ABA and light signalling networks as well as Ca^2+^ influx and K^+^ and Cl^-^ effluxes from SG guard cells (Fig. 6). Interestingly, SG grown plants presented a faster and stronger stomatal recovery when high illumination condition was retrieved, and this was due to the smaller stomatal pore sizes. The faster stomatal response to light may contribute to optimising capsicum photosynthesis under SG light conditions. The current study not only provides valuable physiological implications of SG material on capsicum farming in controlled environment horticulture, but also reveals that SG films could potentially be suitable materials for growing capsicum particularly in the southern hemisphere countries, such as Australia.

**Fig 6.**
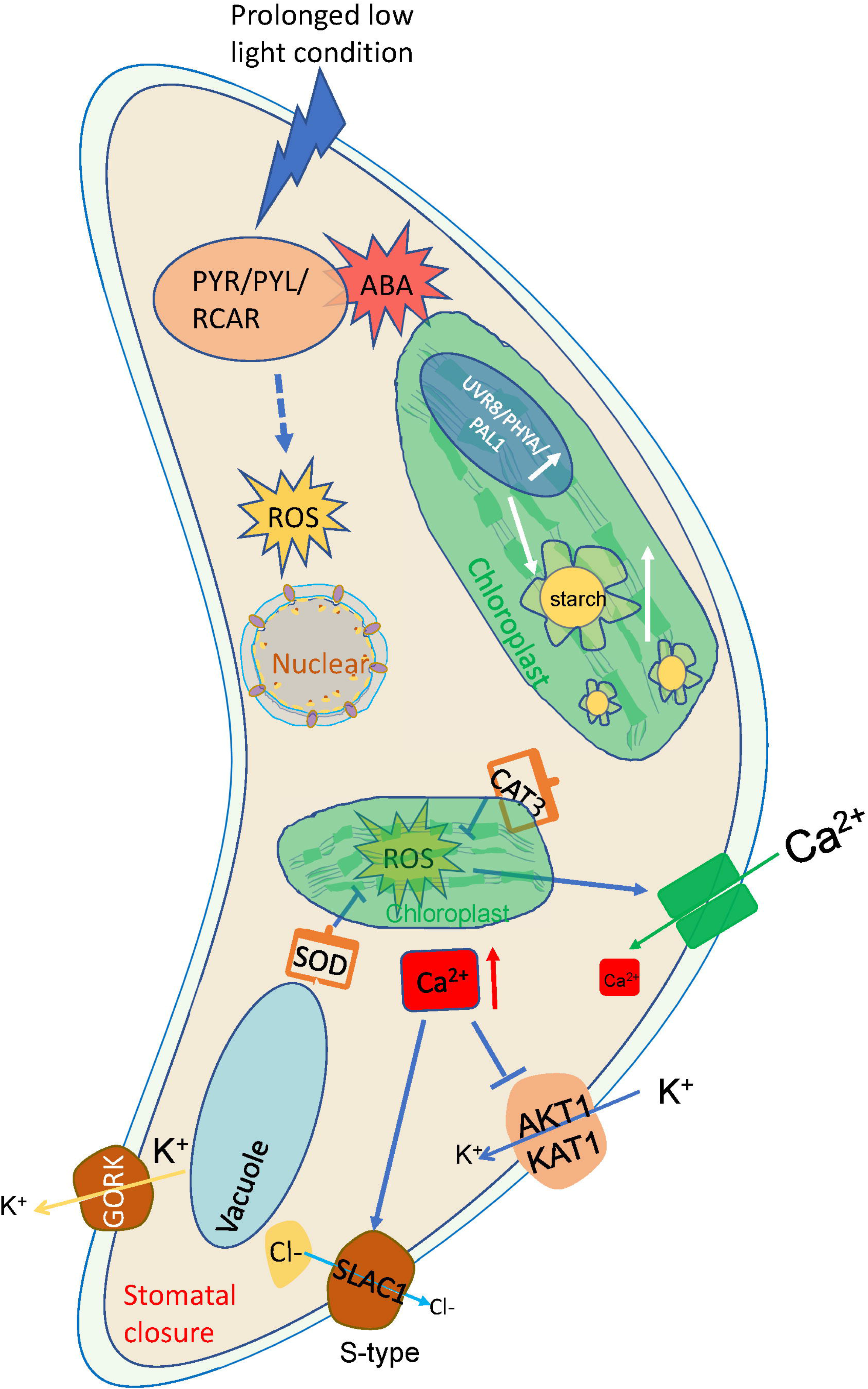
Schematic summary of smart glass (SG)-induced low light condition on guard cell signalling network in capsicum. Under prolonged low light conditions, ABA reception was highly upregulated, leading to the ROS accumulation in guard cells. ROS activated Ca^2+^ influx and thus induced cytosolic Ca^2+^ accumulation in guard cells. Accumulated ROS and Ca^2+^ suppressed K^+^ inward channels AKT1/KAT1 and activated slow anion channels SLAC1(Brandt *et al*., 2012) and K^+^ outward channel GORK. Besides, photosynthesis related genes also play roles in inducing stomatal closure by accumulating starch in guard cells.

## Authors contributions

DT, ZHC, CIC and OG designed the Smart Glass experiment that supported this project. The project was conceived by CZ, ZHC and OG. CZ and SC performed experimental research and data analyses. CZ, ZHC, OG, DT, and CIC wrote the manuscript with contributions from all co-authors.

## Acknowledgments

We thank Dr. Wei Liang for crop growth and management, Ms Chelsea Maier for technical operation and maintenance of the glasshouse, and Dr. Craig Barton for the technical support with Licor 6400XT measurements. We also thank Dr. Juan Zhu from Tasmanian Institute of Agriculture, University of Tasmania for her suggestions on data analysis.

## Supplementary Information

**Fig S1.** Mechanism of smart glass material and an external view of the greenhouse fitted with SG.

**Fig S2.** Effect of smart glass (SG) on net photosynthetic rate of capsicum under light alternations.

**Fig S3.** Correlation analysis of the effect of smart glass (SG) on the maximum theoretical stomatal conductance (*g_max_*), maximum stomatal sizes (*SS_max_*), stomatal opening and closing half-times (*t_1/2_*) with operational stomatal conductance (*g_op_*) of capsicum.

**Fig S4** Relative expression of genes relevant to ABA signalling, ROS metabolism and photosynthesis in capsicum epidermal peels under control and smart glass (SG).

**Table S1** Primers and gene information in the quantitative RT-PCR experiment.

**Table S2.** Comparison of stomatal traits, gas exchange parameters, and ion fluxes of capsicum between control and smart glass (SG) conditions.

